# Uncovering novel endolysins against methicillin-resistant *Staphylococcus aureus* using microbial single-cell genome sequencing

**DOI:** 10.1101/2023.06.22.546026

**Authors:** Takuya Yoda, Ayumi Matsuhashi, Ai Matsushita, Shohei Shibagaki, Yukie Sasakura, Kazuteru Aoki, Masahito Hosokawa, Soichiro Tsuda

**Affiliations:** bitBiome, Inc., 513 Wasedatsurumaki-cho, Shinjuku-ku, Tokyo, 162-0041, Japan; Department of Life Science and Medical Bioscience, Waseda University, 2-2 Wakamatsu-cho, Shinjuku-ku, Tokyo, 162-8480, Japan; Research Organization for Nano and Life Innovation, Waseda University, 513 Wasedatsurumaki-cho, Shinjuku-ku, Tokyo, 162-0041, Japan; Computational Bio Big-Data Open Innovation Laboratory, National Institute of Advanced Industrial Science and Technology, 3-4-1 Okubo, Shinjuku-ku, Tokyo, 169-8555, Japan; Institute for Advanced Research of Biosystem Dynamics, Waseda Research Institute for Science and Engineering, 3-4-1 Okubo, Shinjuku-ku, Tokyo, 169-8555, Japan

## Abstract

Endolysins, peptidoglycan hydrolases derived from bacteriophages (phages), are being developed as a promising alternative to conventional antibiotics. To obtain highly active endolysins, a diverse library of endolysins is vital. We here propose microbial single-cell genome sequencing as an efficient tool to discover dozens of previously unknown endolysins, owing to its culture-independent sequencing method. As a proof-of-concept, we analyzed and recovered endolysin genes within prophage regions of *Staphylococcus* single-amplified genomes (SAGs) in human skin microbiome samples. We constructed a library of chimeric endolysins by shuffling domains of the natural endolysins and performed high-throughput screening against *Staphylococcus aureus*. One of the lead endolysins, bbst1027, exhibited desirable antimicrobial properties such as rapid bactericidal activity, no detectable resistance development, and *in vivo* efficacy. We foresee that this endolysin discovery pipeline is in principle applicable to any bacterial target, and boost the development of novel antimicrobial agents.

## Introduction

The emergence of antimicrobial-resistant bacteria has been a global health concern. Infections with multi-drug resistant bacteria such as ESKAPE pathogens (*Enterococcus faecium, Staphylococcus aureus, Klebsiella pneumoniae, Acinetobacter baumannii, Pseudomonas aeruginosa*, and *Enterobacter* species) are becoming increasingly difficult to treat because of the limited number of antibiotics available.[1] Methicillin-resistant *Staphylococcus aureus* (MRSA) is one of the common pathogens with high mortality. Despite advances in medical research, the management of MRSA infections remains a complex and ongoing clinical challenge, highlighting the urgent need for effective treatment strategies.[2][3]

Endolysins are bacteriophage (phage)-derived enzymes that function as peptidoglycan hydrolases, which can break down the cell wall of host bacteria from within. These enzymes are an essential component of the phage lifecycle, where they help to release new phage particles from the infected host cell. They are considered a promising new class of antimicrobials alternative to conventional antibiotics because of the rapid lysis of bacteria when applied externally.[4][5] Endolysins of *Staphylococcus* phages are mostly composed of two domains, single or multiple enzymatic activity domains (EADs) and a cell-wall binding domain (CBD).[6] Several endolysins have progressed to the clinical trial stage for the treatment of bacterial infections.[7] In addition to their therapeutic potential, endolysins have also found extensive applications in the agriculture and food industries.[8]

Endolysins have several advantages over conventional antibiotics: Their high degree of specificity for the bacterial species or strain has several potential benefits, including a reduced risk of collateral damage to the host microbiota and a lower likelihood of inducing resistance in the target pathogen. Furthermore, owing to the modular composition, protein engineering techniques, such as shuffling modular domains,[9][10] mutagenesis,[11] and conjugation of functional peptides,[12] have been employed to create engineered proteins that have unique and improved characteristics.

In order to fully leverage the potential of endolysins, the availability of a wider range of endolysin genes are desired. The utilization of domain shuffling technique with a library of various endolysins can enhance the therapeutic utility of endolysins. However, a larger pool of endolysin genes is necessary to provide a broader scope of functional domains that can be shuffled to create chimeric proteins with enhanced specificity and activity. Chimeric endolysins are screened in a high-throughput manner,[9][10] and its success depends on the library size and diversity. To obtain endolysin genes, previous research on endolysin development often relied on reference genomes obtained from isolated phages and prophages.[13][14] However, the process of isolation of target bacteria followed by endolysin identification from their genomes are time-consuming and labor-intensive. Recent advancements in metagenomic sequencing have provided access to a vast array of uncultured phage genomes, allowing for a greater understanding of the diversity and unique characteristics of endolysin sequences.[15] Yet, the identification of phage-host associations remains a challenging issue as it requires extensive sequencing data and computing resources.[16]

In this study, we propose that microbial single-cell whole genome sequencing is an efficient method to discover endolysin genes. Previously we developed a hydrogel droplet-based microbial single-cell genome sequencing method called SAG-gel.[17] This method involves the analysis of 96 or more single-amplified genomes (SAGs) and recovers good quality SAGs from various sources, including soil, seawater, and human (fecal, oral, and skin), without relying on culturing of microbial samples.[18–20]

One of the notable features of single-cell genome sequencing is the direct link between endolysin genes and host taxonomic information. In the context of endolysin discovery, this is particularly advantageous as strain-level heterogeneity profiling of mobile genetic elements is much simpler than with metagenomic sequencing. Plasmids and prophages can be easily detected from individual SAGs as demonstrated previously.[20] This feature facilitates the identification of endolysin genes located in prophage regions and allows for immediate association with their target bacteria. Furthermore, identified genes can be recovered by polymerase chain reaction (PCR) from surplus DNA prepared for next generation sequencing, which allows for a low-cost recovery of endolysin genes.

As a proof of concept, we utilized single-cell genome sequencing data derived from human skin swabs to develop novel endolysins against MRSA. A total of 96 endolysins are obtained from 238 SAGs of *Staphylococcus* bacteria together with 29 endolysins from genomes of isolated *Staphylococcus* bacteria. Subsequently, we constructed a chimeric endolysin library by shuffling domains of EAD and CBD. Through this process, we obtained a highly lytic variant bbst1027 against MRSA, which demonstrated desirable antimicrobial characteristics, such as biofilm disruption, sustained activity in human serum, no apparent development of bacterial resistance. Notably, bbst1027 was an artificial endolysin composed of EAD and CBD derived from SAGs. *In vivo* studies using a mouse model infected with MRSA showed that bbst1027 was effective in preventing death due to MRSA infection. Our strategy has the potential to accelerate the development of endolysins by providing a diverse pool of candidates obtained through the culture-independent, single-cell genome sequencing method.

## Results

### Identification of endolysin genes from single-cell amplified genomes

We searched for endolysins against *Staphylococcus aureus* from 675 microbial single-cell genomes from skin swab samples of eight individuals we previously sequenced.[20] We focused on 238 *Staphylococcus* bacterial genomes, including *S. epidermidis, S. aureus, S. capitis, S. hominis, S. saprophyticus, S. lentus, S. warneri, S. sciuri, and S. argenteus*. This approach was chosen due to the similarity in peptidoglycan structures between *Staphylococcus* bacteria. Endolysins discovered from these bacterial genomes are thus likely to be effective against MRSA. Prophage sequences were identified from the genomes, and endolysins were subsequently searched within these prophages by a homology-based method **(Fig. 1A)**. A total of 96 endolysin genes were identified from this analysis. To further expand the endolysin discovery, we sequenced whole genomes of 100 MRSA clinical isolates and 4 food-derived *Staphylococcus* bacteria. The sampling encompassed a diverse range of geographical regions throughout Japan and led to the identification of 29 endolysins.

**Fig 1.**
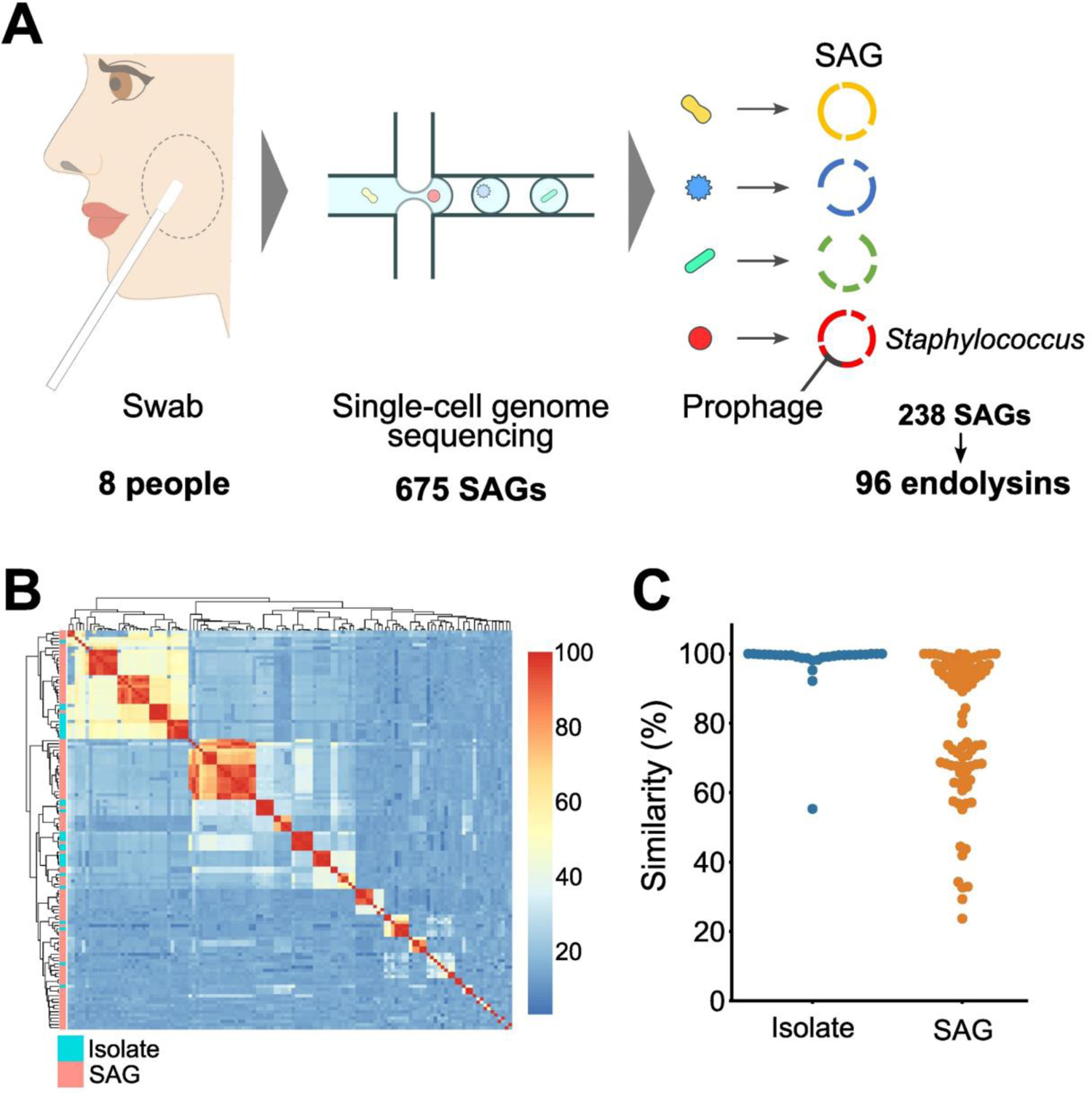
Discovery of endolysins by microbial single-cell genomics. (A) Workflow for the identification of endolysin genes against *Staphylococcus* bacteria from skin swab samples using single-cell genome sequencing. (B) Clustered heatmap analysis comparing amino acid sequences of identified endolysins from isolate genomes and SAGs. (C) Swarm plots depicting similarity distribution between *Staphylococcal* phage lytic proteins in PhaLP and identified endolysin genes from isolate genomes (blue) and SAGs (orange).

We found that the endolysin sequences discovered from SAGs were particularly unique. **Figure 1B** shows the amino acid sequence similarities within the total 125 endolysins. There were notable sequence similarities between endolysins obtained from isolated bacterial genomes (cyan bars), despite the geographical diversity of bacterial samples. In contrast, SAG-derived endolysins collected from eight individuals (red bars) were much more diverse. When compared to the genes in a phage lytic protein database PhaLP,[21] the majority of isolates-derived endolysins showed >95% similarities whereas the SAG-derived ones exhibited significant diversity with similarities as low as 23.7% **(Fig. 1C)**.

### Construction and screening of potent endolysin libraries

Next, we aimed to recover the identified endolysin genes to experimentally demonstrate the potential of unique endolysin genes. From the 125 endolysins, we successfully recovered 96 genes through PCR amplification using either the products of single-cell genome amplification or extracted genomes of cultured samples as templates. However, upon transformation of these endolysin genes into *E. coli*, most of the endolysins exhibited low levels of expression or insolubility. This result was consistent with previous studies demonstrating the difficulty of recombinant production of natural endolysins.[22][23]

To obtain more soluble endolysins, we constructed chimeric endolysins consisting of an EAD and a CBD through domain shuffling. While natural *Staphylococcus* endolysins mostly contain multiple EADs and a CBD, these single EAD-CBD endolysins were expected to be more soluble due to the smaller size. Specifically, we selected 17 EADs (CHAP domains) and 36 CBDs (SH3b and LysM domains) as the building blocks for the chimeric enzymes. CHAP domains were chosen based on the high bactericidal activity in previous studies about engineered endolysins.[10] All the CBD domains identified were adopted for domain shuffling. Homologous overlapping sequences were designed between a vector, EAD, and CBD sequences and introduced by extension PCR (pink, white, and blue squares in **Fig. 2A**). This approach allowed for arbitrary combination of the selected EAD and CBD sequences through Gibson assembly.[24] When transformed into *E. coli*, each colony should contain a chimeric endolysin gene with a unique EAD and CBD pair, potentially generating a library of 576 chimeric endolysins.

**Fig 2.**
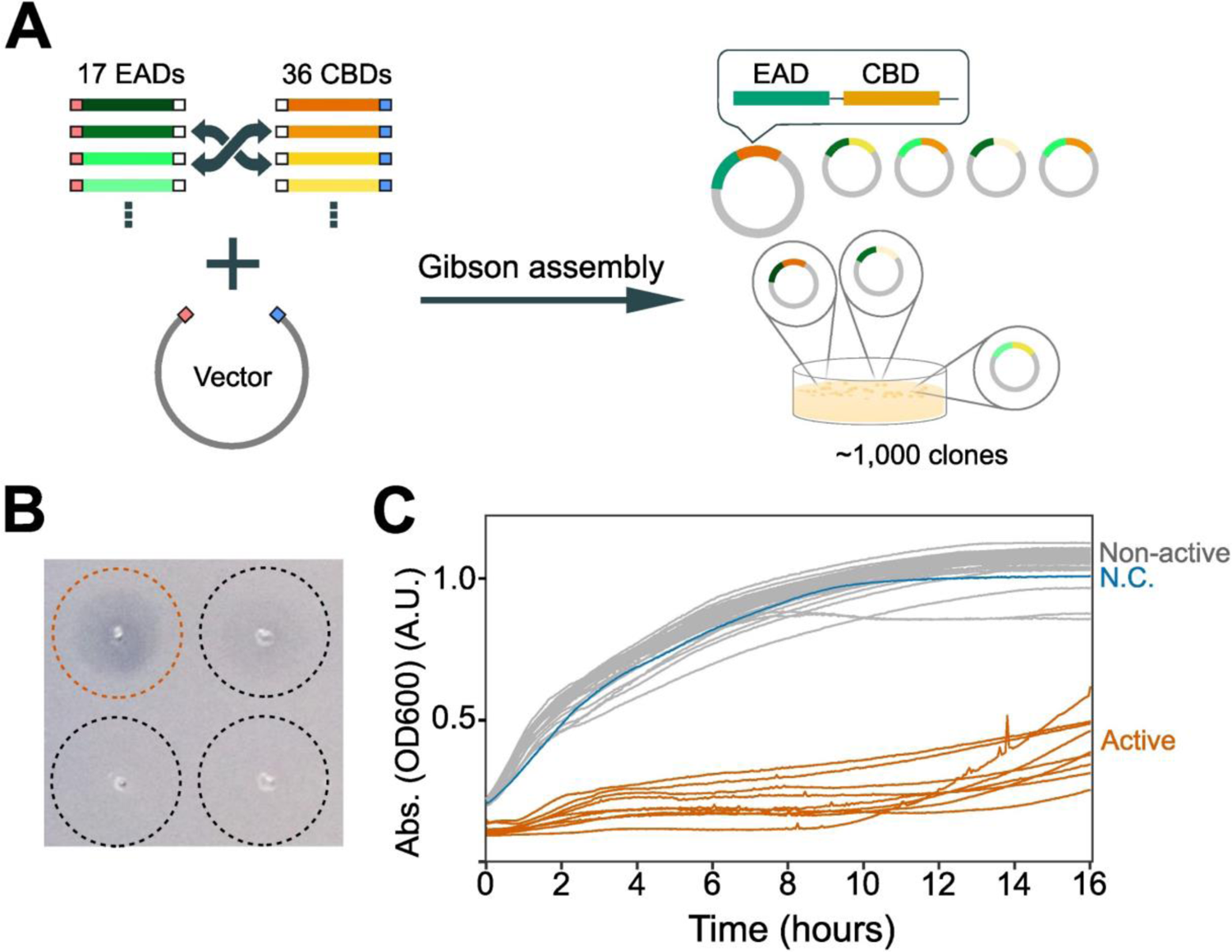
Library construction and screening of chimeric endolysins. (A) A schematic for the construction of a chimeric endolysin library through domain shuffling. One EAD and one CBD are randomly conjugated and cloned into a common vector by Gibson assembly. Each variant is recovered as a colony. (B) The peptidoglycan degradation assay. A dotted circle indicates a spot where an *E. coli* lysate expressing an endolysin was deposited. Enzymatic activity of an endolysin was confirmed by the presence of a clear zone (orange circle). (C) The growth inhibitory assay against MSSA. Lower absorbance values indicate the growth of MSSA was inhibited by active endolysins contained in lysates (orange curves), whereas non-active endolysins exhibited little effect on the growth (gray curves). The negative control (blue curve) indicates the growth curve of MSSA with a lysate that contains an empty vector harboring no endolysin genes.

We screened 746 single clones of the chimeric endolysins using crude lysates to assess the efficacy of engineered endolysins against methicillin-sensitive *S. aureus* (MSSA). The peptidoglycan degradation assay[25] and growth inhibition assay[9] were performed as the first screening step **(Fig. 2B and 2C**). Results of the former assay were visually examined by the presence of lysis plaques with different transparencies. In the growth inhibition assay, samples that exhibited significant decrease in terms of OD600 (0.2 or more units relative to the negative control) were determined active. Samples showed positive results in both assays were identified, totaling 80 variants with a hit rate of 10.7% (80 out of 746). After removing duplicated sequences, a total of 37 variants with unique combinations of EAD and CBD domains were obtained. These lytic endolysins comprised 8 unique EADs from *S. epidermidis, S. aureus, S. capitis, S. hominis*, and *S. saprophyticus*, and 23 unique CBDs from *S. epidermidis, S. aureus, S. capitis*, and *S. hominis*. These results indeed corroborate previous studies that endolysins are efficacious across *Staphylococcus* species[26].

### Identification of engineered endolysin actives against *S. aureus* and its biofilms

The 37 endolysins that exhibited significant peptidoglycan degradation activity were purified, and their activity against *S. aureus* was further evaluated *in vitro*. Although the yields varied depending on the endolysin, all the endolysins were successfully purified (4-10 mg of purified proteins /1L culture). The minimum inhibitory concentration (MIC) of endolysins against the methicillin-sensitive *S. aureus* (MSSA) were determined based on the broth microdilution method (CLSI guideline, M07-A9)[40] and compared to that of vancomycin, a standard of care (SOC) antibiotic for *S. aureus* infection. In comparison at molar concentrations, 28 out of 37 endolysins exhibited a lower MIC than vancomycin, with 8 of them demonstrating remarkable growth inhibition (termed as bbst1001, bbst1005, bbst1014, bbst1021, bbst1027, bbst1035, bbst1036, and bbst1037) **(Fig. S1A)**. The MIC values of these eight endolysins were in the range of 1-4 μg/mL as mass concentrations. The efficacy and composition of these endolysins were shown in **Table S1 and S2**.

Time-kill curve assays were conducted to evaluate the bactericidal activity of the engineered endolysins. The results demonstrated that the turbidity of the MSSA culture decreased significantly within 40 minutes, indicating rapid bactericidal activity. In contrast, vancomycin did not exhibit similar activity under the same conditions, as shown in **(Fig. 3A**. See also **Fig. S1B)**. This result confirms that the mode of action of our endolysins is direct lysis of bacteria, as reported previously.[27] The eight endolysins that demonstrated higher efficacy in the MIC assay were also effective in the time-kill curve assay. Additionally, the eight endolysins we developed were efficient in removing the biomass of *S. aureus* biofilms, while vancomycin and some of the shortlisted endolysins did not exhibit biofilm degrading activity under the same test conditions **(Fig. 3B and Fig. S1C and D)**.

**Fig 3.**
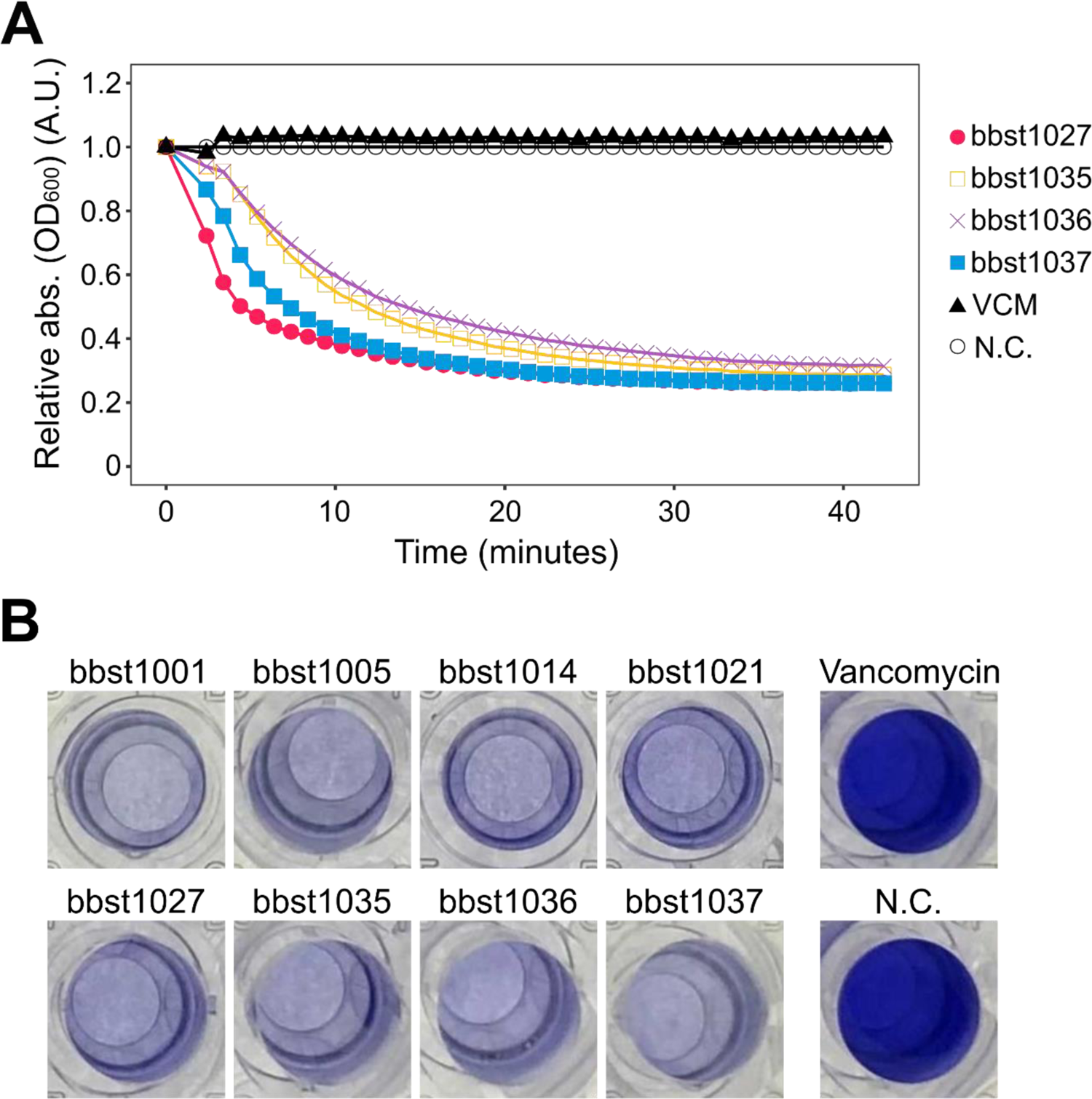
*In vitro* activities of potent endolysin against *S. aureus*. (A) Time-kill curve assay for selected endolysins. Relative absorbance values were calculated by normalizing the absorbance of the negative control at each time point to one. (B) Biofilm degradation assay. The remained biomass in the biofilm of MSSA was stained by crystal violet.

### Efficacy of the endolysins under physiological conditions

To confirm that the endolysins are effective under physiological conditions, we performed following assays:

(1) MIC assays under human serum condition, (2) bacterial spectrum assay, (3) laboratory evolution experiment for the resistance development to endolysins, and (4) *in vivo* animal tests for MRSA infections.

First, we conducted MIC assays in the presence of human serum. We selected a serum concentration of 50% to minimize background noise, as higher serum concentrations had a significant impact on the determination of precise MIC values. Our findings demonstrated that the tested endolysins exhibited equal or improved effectiveness against MSSA in the 50% serum condition as compared to the standard cation-adjusted Mueller Hinton Broth (ca-MHB), shown in **Table 2**.

**Table 2.** Comparison of MIC values of endolysins in the presence or absence of human serum.

|  | MIC (µg/mL) (geometric mean) |  |
| --- | --- | --- |
|  | 50% human serum<br>+ caMHB | caMHB<br>only |
| bbst1001 | 1.6 | 4 |
| bbst1005 | 5 | 6.3 |
| bbst1014 | 2.5 | 1.6 |
| bbst1021 | 1.6 | 5 |
| bbst1027 | 5 | 3.2 |
| bbst1035 | 1.3 | 3.2 |
| bbst1036 | 0.8 | 1.3 |
| bbst1037 | 2.5 | 2.5 |
| Vancomycin | 2.5 | 1 |

We selected 4 endolysins and performed the same assay with nine MRSA clinical isolates **(Table 3**). No significant differences were observed in the MIC values of endolysins between MSSA and MRSA. Notably, the endolysins exhibited activity against some clinical isolates with reduced susceptibility to vancomycin (MIC ≥ 2 μg/mL), as well as those that were sensitive to vancomycin **(Table S3)**.

**Table 3.** Effects of endolysins against clinical isolates of MRSA.

| antimicrobial agent | MIC (µg/mL) |  |  |
| --- | --- | --- | --- |
|  | MIC<sub>50</sub> | MIC<sub>90</sub> | geometric mean |
| bbst1027 | 8 | 8 | 6.3 |
| bbst1035 | 8 | 16 | 7.4 |
| bbst1036 | 2 | 8 | 2 |
| bbst1037 | 8 | 8 | 5.4 |
| vancomycin | 1 | 4 | 1.7 |

The antimicrobial spectrum of our endolysins was also examined using bbst1027 and bbst1037. Growth inhibitions were confirmed for *S. epidermidis* and *S. saprophyticus* as well as *S. aureus* **(Table 4)**. However, they were particularly specific to *S. aureus*. On the other hand, no inhibition of the growth was observed against other Gram-negative and Gram-positive bacteria, which was consistent with previous reports.[10]

**Table 4.** The antibacterial spectrum of endolysins.

|  |  | MIC (µg/mL) |  |  |
| --- | --- | --- | --- | --- |
| Bacteria |  | bbst1027 | bbst1037 | Antibiotics |
| Gram (+) | *Staphylococcus aureus* | 4 | 4 | 1 Vancomycin |
|  | *Staphylococcus epidermidis* | 32 | 32 | 2 Vancomycin |
|  | *Staphylococcus saprophyticus* | 8 | 32 | 2 Vancomycin |
|  | *Streptococcus gallolyticus* | >32 | >32 | 0.5 Vancomycin |
|  | *Bacillus subtilis* | >32 | >32 | 0.25 Vancomycin |
|  | *Enterococcus faecalis* | >32 | >32 | 4 Vancomycin |
| Gram (-) | *Escherichia coli* | >32 | >32 | 0.5 Colistin |
|  | *Acinetobacter baumannii* | >32 | >32 | 2 Kanamycin |
|  | *Pseudomonas aeruginosa* | >32 | >32 | 0.5 Colistin |

The MIC values of endolysins against various bacteria were shown.

Next, we conducted laboratory evolution experiments by culturing bacterial population under endolysin stress for 11 days. The MSSA strain NBRC 100910 was serially passaged in the presence of sub-MIC endolysins or vancomycin for 11 days. Unexpectedly, the MIC values of most endolysins except for bbst1027 increased more than eight-fold in a relatively short term, while those of bbst1027 remained mostly constant (up to two-fold, **Fig. 4A**). This would be due to the most rapid bactericidal activities of this endolysin among the endolysins we obtained, as we shown in **Fig. 3A**. Indeed, the treatment with bbst1027 at 1×MIC concentration demonstrated more than 5 log reductions of *S. aureus* within 30 minutes and did not show the regrowth of bacteria even after 24 hours, whereas bbst1037 showed less than 2 log reductions within 1 hour and the regrowth of bacteria was observed afterwards **(Fig. 4B**). Comparing bbst1027 with bbst1037, we concluded that bbst1027 was more potent than bbst1037 because significant bactericidal activities were observed even at the lower concentrations **(Fig. 4C**). We speculated that this was partly due to the positively charged EAD of bbst1027 **(Fig. 4D**). As suggested previously,[28][29] the positive charge of endolysin contributes to accessibility to the bacterial cell surface. Both EAD and CBD of bbst1027 were positively charged, whereas only CBD was positively charged in other endolysins.

**Fig 4.**
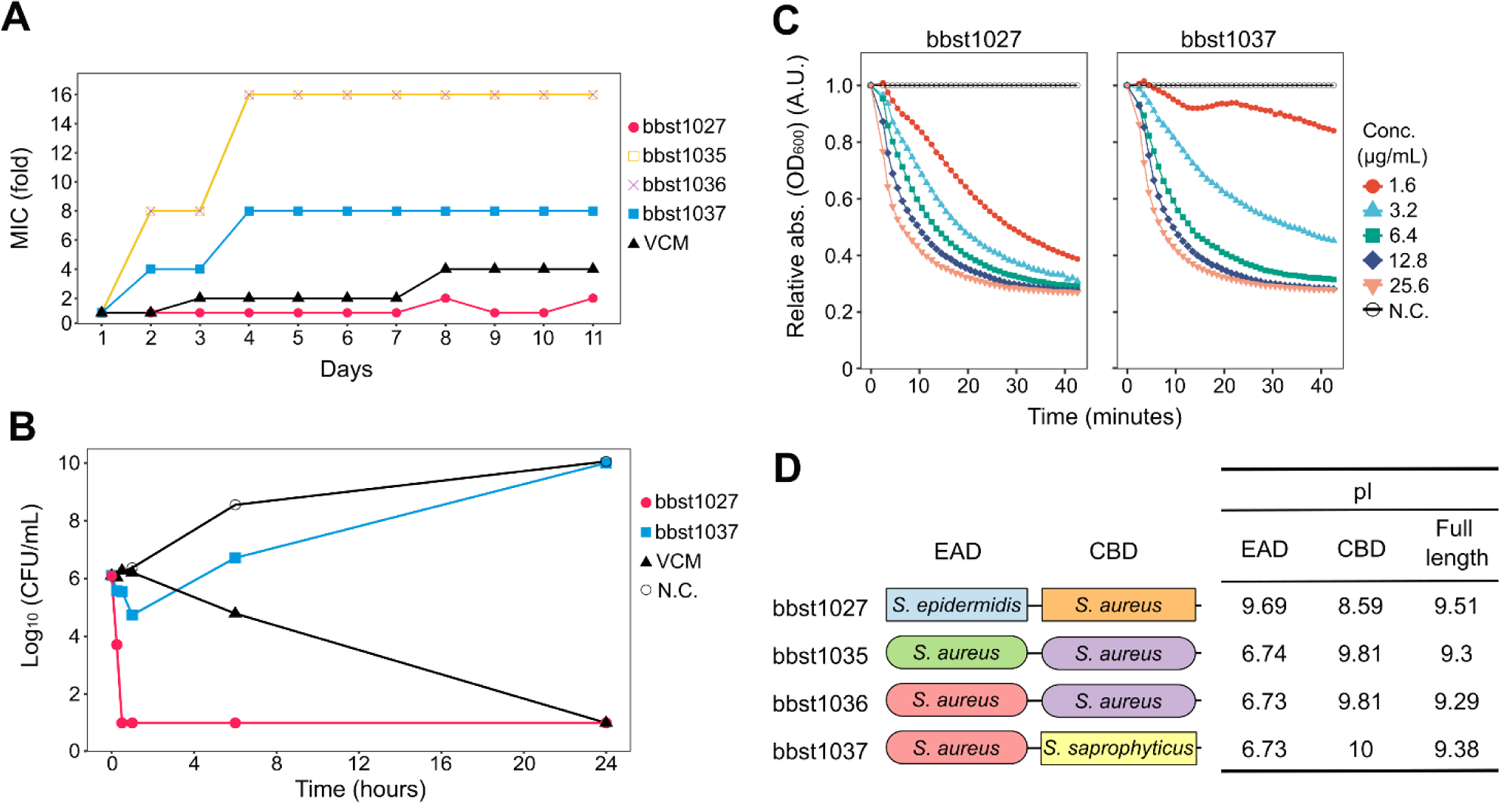
In vitro activities of bbst1027. (A) Laboratory evolution experiment of MSSA under antimicrobial stress for 11 days. (B) Time-kill curve of 1xMIC endolysins or vancomycin against MSSA. The colony forming units (CFU) at each time point were shown with 10 CFU/mL as the limit of detection for this assay. (C) Dose responses of bbst1027 and bbst1037. Relative absorbance values were calculated by normalizing the absorbance of the negative control at each time point to one. (D) Schematic structures and isoelectric point (pI) values of endolysins and their domains. The shapes of endolysin domains indicate the sample origins. Square and oval shapes represent domains derived from SAGs and isolated genomes.

The above *in vitro* findings indicate that bbst1027 has potential therapeutic efficacy against MRSA infections. After confirming the absence of hemolytic and cytotoxic activities of bbst1027 (**Fig. S2**), we evaluated *in vivo* efficacy of bbst1027 using the MRSA-induced bacteremia mouse model. The clinical isolate MRSA No. 76 used in this assay was resistant to oxacillin but sensitive to vancomycin and exhibited hemolytic activity on 5% sheep blood agar plates. The MIC value of bbst1027 to this clinical isolate was 8 μg/mL **(Table S3)**. Upon intravenous injection of MRSA No. 76 at doses of 1×10^7^ CFU/head or 2×10^7^ CFU/head, all mice succumbed to infection within 24 hours, with the higher dose showing more severe symptoms (**Fig. 5**). However, three injections of 20 mg/kg of bbst1027 significantly improved the survival rate of bacteremic mice in both doses of MRSA (p < 0.001 in the 1×10^7^ CFU/head condition, p = 0.002 in the 2×10^7^ CFU/head condition). These results demonstrate the *in vivo* efficacy of bbst1027 as a potential therapeutic for *S. aureus* infection.

**Fig 5.**
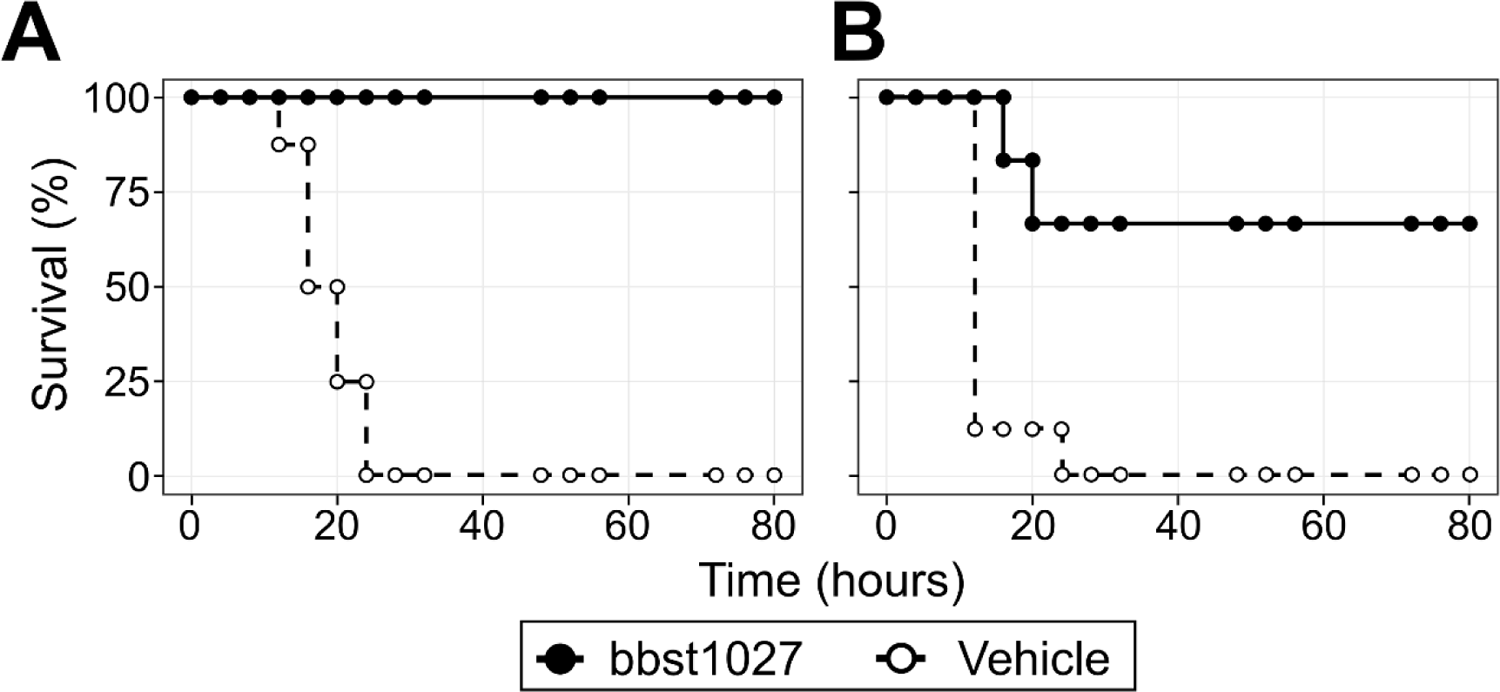
*In vivo* efficacy of bbst1027 in MRSA-induced mouse bacteremia model. BALB/c mice were intraperitoneally administered the MRSA clinical isolate No. 76 at (A) 1×10^7^ CFU/head and (B) 2×10^7^ CFU/head. Mice were then intraperitoneally administered 20 mg/kg of bbst1027 (n=6) or vehicle (n=8) at 1, 3, and 5 hours after MRSA inoculation. P-values in bbst1027 treatment were <0.001 (A) and 0.002 (B) at the log-rank test.

## Discussion

In this study, we demonstrated that microbial single-cell genome sequencing was an effective method to obtain endolysin genes from bacterial genomes. This approach allows for direct association between a bacterial host and its genes, which is a key feature for efficient discovery endolysins targeting a specific bacterium. Notably, our single-cell sequencing method required only eight skin swab samples to yield approximately 100 unique *Staphylococcus* endolysin genes. These genes were readily recoverable from amplified genome DNA through a simple PCR procedure, circumventing the need for costly DNA synthesis.

Our method allows a facile construction of diverse endolysin library, which provides better starting point for highly effective antimicrobials. Endolysin sequences acquired from single-cell genome data contained low homology sequences (39 out of 96 genes with homology less than 80%) compared to the public endolysin database PhaLP. Despite the low sequence homology, they were confirmed to be functional as endolysins. The EAD domain of bbst1027 developed in this study showed 69% sequence similarity to the closest sequence in the PhaLP database. By contrast, most of the clinical isolates matched the known endolysins (all but one had more than 90% homology), even though they were obtained from various locations in Japan. These findings suggest that culture-based identification methods might introduce inherent biases and may overlook a diverse set of endolysin genes that cannot be accessed through conventional methods.

Unique endolysins enable the creation of an even larger engineered endolysin library through various protein engineering techniques, including the domain shuffling method employed in this study. As demonstrated in previous studies,[9][10] domain shuffling is particularly well-suited to the modular nature of endolysin genes, making it possible to develop highly active engineered endolysins that can target not only Gram-positive but also Gram-negative bacteria.

To efficiently explore the vast potential of endolysins as antimicrobials, it is essential to implement screening methods that can handle the screening of a large endolysin library. While the domain shuffling method utilized in this study allows for the generation of an expanded library of engineered endolysins, it is important to note that the identification of the most effective candidates is labor-intensive and requires efficient yet rigorous screening. We speculate that new technologies, such as *in silico* modeling using AlphaFold2[30] and microfluidic *in vitro* ultra-high-throughput screening [31] can be employed in a near future to expedite the screening process and enhance the identification of highly potent endolysins within the large library.

While the medical applications of endolysins are well-known, there are numerous other potential applications for these enzymes[32]. In particular, the ability of endolysins to degrade biofilms could have significant applications in various industries. Biofilms are ubiquitous and can form on a wide range of surfaces, including medical devices, food processing equipment, and even ship hulls.[33] For example, *Pseudomonas aeruginosa* biofilms are known to be problematic in cystic fibrosis patients, and endolysins can be used to break down these biofilms and prevent further infection. In addition to medical applications, endolysins could be used in the food industry to prevent biofilm formation and contamination of food processing equipment.[34] Biofouling and biocorrosion are also significant problems in various industrial settings, and endolysins could be used to prevent the buildup of biofilms on pipes, tanks, and other equipment. By degrading biofilms, endolysins could help reduce the need for harsh chemical cleaners and disinfectants, making them a potentially more environmentally friendly solution. Most biofilms consist of a mixture of different bacteria, making it challenging to identify and target specific species for elimination. The microbial single-cell genome sequencing method offers a solution to not only identify the detailed taxonomic information of biofilm-forming bacteria, but also discover endolysins from their genomes, which can be used to eliminate them.

## Conclusion

In conclusion, our study highlights the potential of microbial single-cell genome sequencing as a powerful tool for discovering novel endolysins and constructing an engineered endolysin library. Our lead endolysin, bbst1027, showed strong activity against MRSA and demonstrated several favorable characteristics such as rapid lysis, narrow antibacterial spectrum, biofilm elimination, and low levels of resistance development. Its ability to target biofilms and remove biomass makes it a promising candidate for combination therapy with antibiotics and treatment of infected implants. Moreover, bbst1027 was effective in a mouse model and did not exhibit hemolytic or cytotoxic effects, suggesting its potential for clinical applications in the future. Further studies are needed to optimize its conditions for clinical trials, but the findings of our study provide a solid foundation for the development of endolysin-based therapeutics against MRSA infections and other bacterial pathogens.

## Materials and methods

### Bacterial strains, media, antibiotics, and growth conditions

The bacterial strains used in this study are as follows; methicillin-sensitive *Staphylococcus aureus* (MSSA) strains NBRC 100910 and NBRC 13276 (National Institute of Technology and Evaluation, NITE), *Staphylococcus epidermidis* strain NBRC 113847 (NITE), *Staphylococcus saprophyticus* strain NBRC 102446 (NITE), *Streptococcus gallolyticus* strain ATCC BAA-2069 (BEI resources), *Bacillus subtilis* strain ATCC6633, *Enterococcus faecalis* strain NBRC 100482 (NITE), *Escherichia coli* K12 strain ATCC 10798, *Acinetobacter baumannii* strain NBRC 110489 (NITE), *Pseudomonas aeruginosa* strain NBRC 12582 (NITE). Clinical isolates of methicillin-resistant *S. aureus* were obtained from Gunma University Graduate School of Medicine. Food-derived *Staphylococcus* bacteria were isolated from chicken and pork obtained from supermarket stores in Japan. The bacterial strains were grown in the following liquid media at 37 °C unless otherwise mentioned; tryptic soy broth (TSB) with 7.5% NaCl (Nacalai Tesque) for *S. aureus*, TSB (Becton Dickinson, BD) for *S. epidermidis* and *S. gallolyticus*, No.802 medium (Hipolypepton 10g, Yeast extract 2g, MgSO_4_·7H_2_O 1g in 1L distilled water) for *S. saprophyticus*, LB broth, Miller (Nacalai tesque) for *A. baumannii, E. coli* and *P. aeruginosa*, brain heart infusion medium (Sigma-Aldrich) for *B. subtilis*, and Lactobacilli MRS broth (BD) for *E. faecalis*. Antibiotics used were all obtained from Fujifilm Wako Pure Chemical, except daptomycin from Tokyo Chemical Industry.

### DNA extraction and genome sequencing

DNAs of the 100 clinical isolates of *S. aureus* and 4 food-derived *Staphylococcus* bacteria were extracted using the DNeasy 96 Blood & Tissue kit (Qiagen). For sequencing analysis, sequencing libraries were prepared using the QIAseq FX DNA Library Kit (Qiagen). Each library was sequenced using the Illumina NextSeq2000 2 × 150 bp configuration.

### Identification of endolysin genes from microbial genome data

Single-cell genome sequencing data of *Staphylococcus* bacteria used in this study was obtained in our previous study.[20] Prophage regions in host bacterial genomes were identified by PhageBoost,[35] and annotated by PHANOTATE.[36] Endolysin genes were searched by DIAMOND[37] using endolysin genes in the RefSeq database as reference. Pfam domains in the endolysin genes were identified by InterProScan.[38] Endolysin candidates were selected based on the presence of Pfam domains. Endolysin candidates were selected based on the preference of Pfam domains previously reported to be associated with endolysin genes in the literature.[39]

### Construction of the natural endolysin library

Endolysin genes were amplified by PCR using the products of single-cell genome amplification or extracted genomes of cultured samples as templates. PCR products were purified using the Wizard SV Gel and PCR Clean-Up System kit (Promega) and ligated into pET17b vector using the Gibson Assembly Master Mix (New England Biolabs). The ligation products were transformed into *E. coli* BL21(DE3)pLysS (Promega) on LB agar plates containing 100 μg/mL ampicillin.

### Construction of the artificial endolysin library

Domains annotated either as cysteine, histidine-dependent amidohydrolases/peptidase (CHAP), SRC Homology 3 (SH3), or Lysin Motif (LysM) were individually amplified by extension PCR to attach homologous overlapping sequences using the endolysin genes obtained above. The PCR products of 17 EADs and 36 CBDs were generated in total with common homologous sequences on both ends for EAD and CBD (see **Fig. 2A**). EAD and CBD amplicons were mixed at an equal molar ratio, and ligated into the pET17b plasmid randomly using the Gibson Assembly Master Mix (NEB). The transformation was performed in the same manner as the above.

### Endolysin expression and lysate preparation for high-throughput screening

Lysates were prepared by the modified protocol of Gerstmans et al., 2020.[9] Briefly, Single colonies were picked into wells of a 96-deep-well plate filled with 500 μl of LB with ampicillin (100 μg/ml) and incubated overnight at 37 °C. Twenty-five microliters of cultures from each well were transferred to a new 96-deep-well plate, each filled with 500 μL of the Overnight Express Autoinduction System 1 (Novagen) per well and incubated at 37 °C for 5 hours, followed by incubation overnight at 16 °C. After the collection of cells by centrifugation, the pellets in a 96-deep-well plate were treated with chloroform vapor for an hour at room temperature. To remove the residue chloroform, the plate was incubated at 37 °C for 30 minutes. Each pellet was then suspended in 500 μL lysis buffer (20 mM HEPES-NaOH (pH 7.4), 150 mM NaCl), added 0.2U of DNase I (Takara), and incubated at 30 °C for an hour. Finally, lysates were centrifuged at 2,000g at 4 °C for 5 minutes and stored at 4 °C until use.

### Peptidoglycan-degradation assay

*S. aureus* (NBRC 100910) cells were grown to the stationary phase in 100 mL of Tryptic Soybean Broth (TSB). The cultures were autoclaved and harvested at 6,000g for 10 minutes. The pellet was suspended in 3 mL of lysis buffer and added to 150 mL of lysis buffer with 0.8% agarose at 60 °C. The mixture was poured into a one-well plate (Stem). After setting the agarose gel, 5 μL of the cell lysate was spotted onto the agar plate and incubated overnight at 37 °C. The degradation activities were evaluated by the appearance of clear zones after incubation for 1 hour.

### Growth inhibition assay

*S. aureus* (NBRC 100910) cells were grown overnight at 37 °C in TSB with 7.5% NaCl. The cultures were diluted to OD600 = 0.1 with TSB with 7.5% NaCl. The diluted cultures and the lysate were mixed in equal amounts (50 μL) and incubated at 37 °C for 16 hours. The growth of the cells were monitored by measuring at OD600 using Infinite 200 PRO M Nano+ (Tecan). The criterion for assessing activity involved quantifying the OD600 of each well on the plate after a 16-hour incubation period. Samples exhibiting a difference of 0.2 or greater in OD600, compared to the measurement obtained from wells where PBS was added instead of lysate as a negative control, were classified as exhibiting significant activity.

### Expression and purification of endolysins

To express the endolysins, the BL21(DE3)pLysS strain carrying the endolysin gene-cloned pET17b plasmids was grown in LB broth until the OD600 reached 0.4-0.8. The bacterial cultures were then induced with 0.5 mM IPTG (Nacalai Tesque) overnight at 16 °C. Bacterial cells were harvested and suspended with xTractor™ buffer (Clontech) and then treated with appropriate amounts of proteinase inhibitors (Takara Bio) and DNase I (Takara Bio) on ice until the bacterial cell are lysed. The supernatant was obtained by centrifugation of the cell lysate at 10,000 *g* for 30 minutes and applied into the His TALON™ gravity column (Clontech). The columns were washed to remove the non-target proteins, and the target proteins were eluted with phosphate-buffered saline (PBS) (-) (pH 7.4) containing 150 mM imidazole. The eluates were dialyzed with PBS(-) (pH 7.4). For *in vivo* use, the purified endolysins were obtained by purifying the supernatant with a linear gradient of 10-150 mM imidazole using the HisTrap HP column (Cytiva). The eluates were concentrated with Amicon Ultra-15, 3k MW cut-off (Millipore) if necessary. After dialysis with PBS, the contaminated endotoxin was removed with Pierce™ High Capacity Endotoxin Removal Resin (Pierce).

### Minimum inhibitory concentration (MIC) assay

MICs of endolysins and antibiotics were determined by the broth microdilution methods[40] based on CLSI guideline M07-A9. The overnight culture of bacteria was inoculated into the designated growth medium and was cultivated for approximately 4 hours at 37 °C. The bacterial cells were harvested and suspended in cation-adjusted Mueller-Hinton broth (ca-MHB). The cells were adjusted to 5×10^5^ colony-forming units (CFU)/mL and exposed to endolysins or antibiotics in a series of two-fold serial dilutions in the 96-well microtiter plate. After 18 hours of incubation at 37 °C, MIC values were determined. In this study, the MIC was defined as the lowest drug concentration that suppressed bacterial growth below OD600 < 0.2. We confirmed that the MIC values determined like this were almost identical to those by the visual judgment as defined.

The MIC values in human serum were determined as described above, with one exception. Specifically, 50% human serum (KAC), in which the complement components were not inactivated, was added to the reaction mixture. Due to increased background interference in this condition, the MIC values were determined by visual inspection in accordance with CLSI guidelines since it was not possible to determine the MIC value by measuring optical density.

### Minimum bactericidal concentration (MBC) assay

MBC values were determined according to the CLSI guideline[41] (CLSI guideline M26A), with procedures similar to those used for MIC determination as described above. After determining the MIC values, 10 μL of reaction mixtures corresponding to sub-MIC, MIC, and above MIC were spread onto TSB agar plates containing 7.5% NaCl. All agar plates were incubated at 37 °C overnight, and the MBC values were determined in accordance with the CLSI guideline M26A.

### Time-kill assay

Time-kill curve was evaluated using two different ways; the consecutive measurement of turbidity of the bacterial cell suspensions for short-term (40 minutes) or the CFU measurement at the designated time points up to 24 hours. For the former method, overnight culture of bacterial cells was centrifuged, and the cell pellets were resuspended into the reaction buffer (20 mM Tris pH 7.5, 150 mM NaCl, 0.9 mM MgCl_2_, 2.5 mM CaCl_2_). This cell suspension was adjusted to OD600 = 0.7 with the reaction buffer, and then 1/10 volume of endolysins was added at the concentration of 25.6 μg/mL. The OD600 was immediately measured and kept being monitored for 40 minutes at room temperature. For the latter method, bacterial cells of the overnight culture were harvested and suspended into ca-MHB to adjust the bacterial concentration to 5×10^5^ CFU/mL. The cells were then exposed to endolysins and antibiotics at their respective MIC values and incubated at 37 °C. At designated time points, a portion of the culture was sampled and diluted over 100-fold before spreading onto TSB agar plates with 7.5% NaCl. The agar plates were incubated overnight at 37 °C and the CFU was determined.

### Biofilm assays

The process of *S. aureus* biofilm formation followed the method described elsewhere.[42] An overnight culture of *S. aureus* NBRC 13276 was adjusted to OD600 = 0.1 and diluted 100-fold with TSB containing 0.2% glucose (TSBg) before being inoculated into a flat-bottom 96-well polystyrene plate (VIOLAMO). The plate was incubated at 37 °C for 24 hours to facilitate biofilm formation, after which the planktonic cells were removed by washing with PBS three times. Endolysins and antibiotics in PBS were added to the plate and incubated at 37 °C for 24 hours. Biofilm disruption was assessed using the crystal violet stain method [42]. Following a 24-hour exposure to antimicrobial agents, detached bacterial cells were removed by washing with phosphate-buffered saline (PBS) twice. Subsequently, the remaining attached bacteria were stained with crystal violet for 15 minutes, followed by removal of the supernatant and further washing with PBS. The crystal violet bound to the bacteria was resolubilized by adding 33% acetic acid, and the OD590 of the supernatants from each well was measured.

### Laboratory evolution experiment for resistance development to endolysins and antibiotics

The laboratory evolution experiment was carried out as a serial passage MIC experiment in the presence of antimicrobial agents at the sub-MIC concentration. *S. aureus* NBRC 100910 susceptible to all the test compounds and antibiotics was used. At each step, the MIC was measured as described above, then a cell culture corresponding to the sub-MIC concentration was transferred to another round of MIC experiment. The inoculums were adjusted to OD_600 nm_ = 0.1 and further diluted 100-fold with ca-MHB. The final solution contained approximately 1×10^6^ CFU/mL of bacterial cells. The dilutions were prepared for each serial experiment. The process was repeated for 11 days and bacteria grown at the sub-MIC concentrations were stored at −80 °C in caMHB with 16% glycerol each day. For the final MIC analysis, the glycerol stock samples were passaged twice in the absence of any antimicrobial compounds and MICs were measured as described above.

### Mouse bacteremia model

The MRSA clinical isolate No. 76, obtained from Gunma University, was suspended in saline containing 5% mucin to a concentration of 5×10^7^ CFU/mL. Mice were then intraperitoneally injected with 0.2 mL of the bacterial cell inoculum (1×10^7^ CFU/head), followed by intraperitoneal administration of the vehicle (PBS) and test compounds at 1, 3, and 5 hours after MRSA inoculation. Endolysins and vehicle were administered via the intraperitoneal route, while vancomycin was administered subcutaneously. Clinical symptoms were monitored for 3 days post-MRSA inoculation, and the survival rate was analyzed statistically using the log-rank test.

## Data availability

The assembled genomes of 4 clinical isolates of S. aureus and 4 food-derived Staphylococcus bacteria were deposited in DDBJ/ENA/GenBank under BioProject number PRJNA981294.

