## Supporting information for "Uncovering novel endolysins against methicillin-resistant *Staphylococcus aureus* using microbial single-cell genome sequencing"

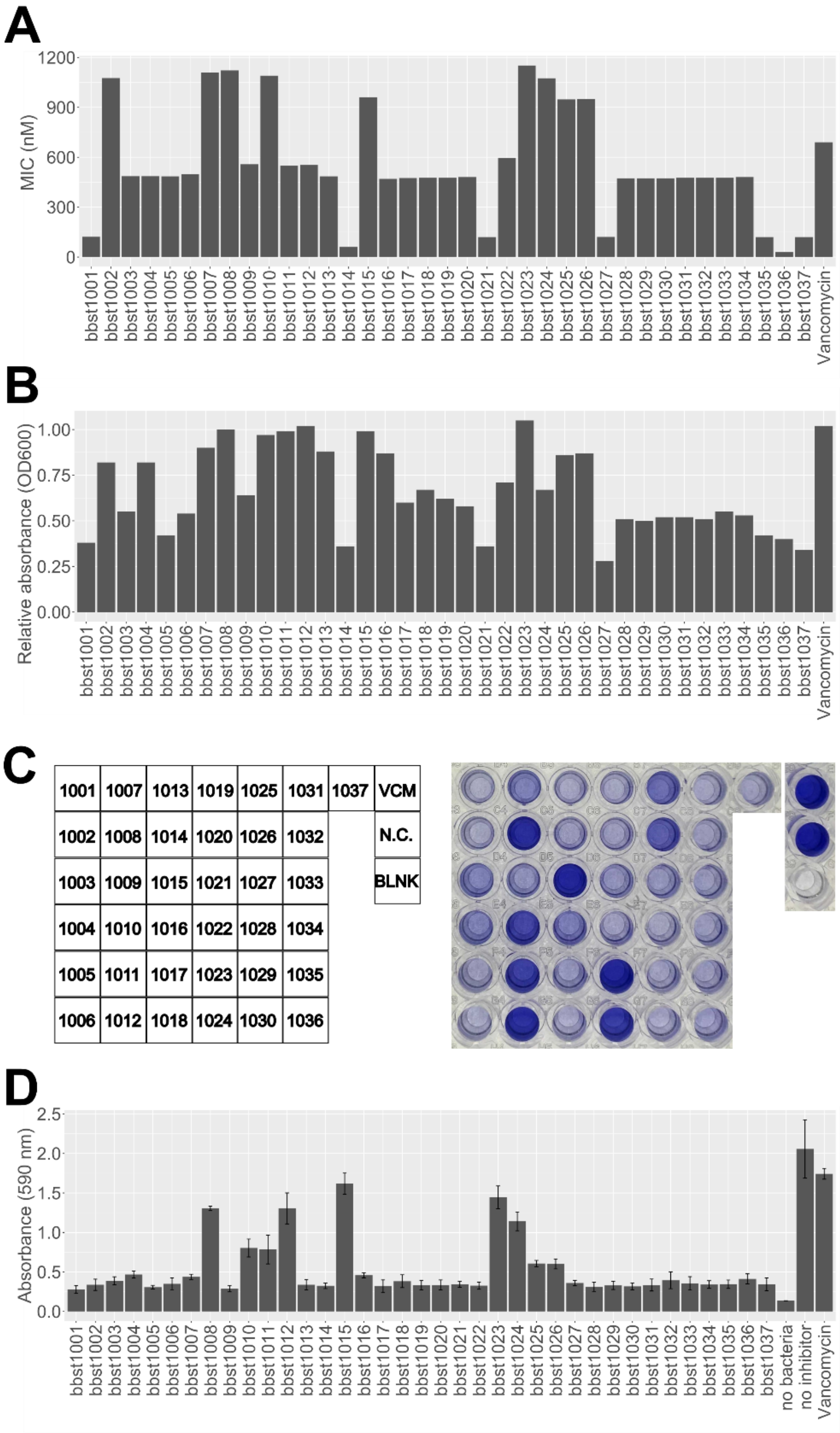


**Fig. S1 *In vitro* activities of endolysin against S. aureus.** (A) MIC values of endolysins were determined. The MIC values of endolysins against S. aureus strain NBRC 100910 were shown as a molar concentration. The bbst1001~bbst1037 were clone names of individual engineered endolysins we newly generated. (B) The bactericidal activities of endolysins were shown. S. aureus strain NBRC 100910 was treated with 25.6 µg/mL of endolysins or vancomycin at 25 °C, and absorbance at OD600 of cultures was monitored for 40 minutes. The relative absorbance at OD600 was calculated by normalization of the OD600 values at the endpoint, namely at 40 minutes, by the values of the growth control. (C) Biofilm destruction by endolysins. Biofilm of S. aureus strain NBRC 13276 was treated with each endolysin and vancomycin in triplicates. (D) The remained biomass of biofilm after treatment was stained by crystal violet and was detected by measurement of the optical density at 590 nm.

**Table S1 Characterization of 8 hits selected after the first screening campaign.**

| ID | Sequence | | Similarity to PhaLP(%) | |
|---|---|---|---|---|
| | EAD | CBD | EAD | CBD |
| bbst1001 | MQAKLTKKEFIEWLKTSEGKQYNADG<br>WYGFQCFDYANAGWQVLFGYNLKGV<br>GAKDIPSANDFNGLATVYQNTPDFLAQ<br>PGDMVVFGSNYGAGYGHVAWVIEAT<br>LDYIIVYEQNWLGGGWTDGVQQPGSG<br>WEKVTRRQHAYDFPMWFIRPNFKSET<br>APRSVQSPTQASKKET | **EIAKQEVLPTGWKKNKHGT<br>YYKAQKGSFINGNQPIQARY<br>VGPFRLKNNAAGDLPANTK<br>IEYDEIMLQDKHVWVGYDS<br>FEGERIYLPVGTWNGKKPP<br>KNKMKQVWGIL** | 100 | **47.959** |
| bbst1005 | MQAKLTKKEFIEWLKTSEGKQYNADG<br>WYGFQCFDYANAGWQVLFGYNLKGV<br>GAKDIPSANDFNGLATVYQNTPDFLAQ<br>PGDMVVFGSNYGAGYGHVAWVIEAT<br>LDYIIVYEQNWLGGGWTDGVQQPGSG<br>WEKVTRRQHAYDFPMWFIRPNFKSET<br>APRSVQSPTQASKKET | **VKNKPGSASTPANRRDMSG<br>WKINKYGTYYKSEVAHFTP<br>NTPIKTHYVGPFRSCPVSGV<br>LQPGQIVRYDTVCKQDGHV<br>WISYTAYNGKDVWLAVRT<br>WDKNTDSLGKLWGTIN** | 100 | 97.273 |
| bbst1014 | MQAKLTKKEFIEWLKTSEGKQYNADG<br>WYGFQCFDYANAGWQVLFGYNLKGV<br>GAKDIPSANDFNGLATVYQNTPDFLAQ<br>PGDMVVFGSNYGAGYGHVAWVIEAT<br>LDYIIVYEQNWLGGGWTDGVQQPGSG<br>WEKVTRRQHAYDFPMWFIRPNFKSET<br>APRSVQSPTQASKKET | **VKNKPGSASTPANRRDMNG<br>WKINKYGTYYKSEVARFTP<br>NTPIKTHYVGPFRSCPVSGV<br>LQPGQTIKYDTVCKQDGHV<br>WVSYTAYNGKDVWLAVRT<br>WNKTNDSLGKLWGTIN** | 100 | 97.297 |
| bbst1021 | **MKTQSQINARLNAYKNGTVDSPYRV<br>KTWTSYDPAFGTMEPGCIDVDHAYH<br>AQCADLPIDYILWLTDNQYRAWGNA<br>KDFPNNKFPTGWKVIENLPSTVPQK<br>GWIAVFSSGTYAQYGHIGLVYDGGN<br>TNSFEILEQNWNGYANKKPTLRWDN<br>YYGLTHFIVPPVAKEVHTLTTKVKE<br>APKQT** | **TKKTSTSNEKWNKNQYGIL<br>WRKEVGSFTCNVPQGIITRR<br>IGPGRQYPIAGALKKAQTV<br>NYTEIQKNDGYIWISWMTN<br>SGYTVYMPVRQVKSDGSLG<br>PLWGTIK** | **73.889** | **65.591** |
| bbst1027 | **MAKTQTQINKLVDSYLGKYVDFDGY<br>YAFQCMDLAVSYVYKLTDGSFRMY<br>GNAKDAINNKFPSGWKVIRNQAATV<br>PKKGWIAVYTTGVYQQYGHIGIVYN<br>GGNTSQFQILEQNFDGLANSPAKLR<br>WDNYSGLTHFIVPPTKSATTSSNGSA<br>KTTTAKAKTSTPKTKKRKIMLVAGH<br>GYNDP** | **SSNTVKPVASAWKRNIYGT<br>YYMEESARFTNGNQPITVR<br>KVGPFLSCPVGYQFQPGGY<br>CDYTEVMLQDGHVWVGYT<br>WEGQRYYLPIRTWNGSAPP<br>NQILGDLWGEI** | **67.614** | 97.143 |
| bbst1035 | MQAKLTKKEFIEWLKTSEGKQFNVDL<br>WYGFQCFDYANAGWKVLFGLLLKGL<br>GAKDIPFANNFDGLATVYQNTPDFLAQ<br>PGDMVVFGSNYGAGYGHVAWVIEAT<br>LDYIIVYEQNWLGGGWTDRIEQPGWG<br>WEKVTRRQHAYDFPMWFIRPNFKSET<br>APRSIQSPTQASKKETAKPQPKAVE | KNQKNPPVPAGYTLDKNNVP<br>YKKETGNYTVANVKGNNVR<br>DGYSTNSRITGVLPNNATIKY<br>DGAYCINGYRWITYIANSGQR<br>RYIATGEVDKAGNRISSFGKF<br>STI | 100 | 100 |
| bbst1036 | MQAKLTKKEFIEWLKTSEGKQFNVDL<br>WYGFQCFDYANAGWKVLFGLLLKGL<br>GAKDIPFANNFDGLATVYQNTPDFLAK<br>PGDMVVFGSNYGAGYGHVAWVIEAT<br>LDYIIVYEQNWLGGGWTDGIEQPGWG<br>WEKVTRRQHAYDFPMWFIRPNFKSET<br>APRSVQSPTQAPKKETAKPQPKAVE | KNQKNPPVPAGYTLDKNNVP<br>YKKETGNYTVANVKGNNVR<br>DGYSTNSRITGVLPNNATIKY<br>DGAYCINGYRWITYIANSGQR<br>RYIATGEVDKAGNRISSFGKF<br>STI | 99.444 | 100 |

| | | | | |
|---|---|---|---|---|
| bbst1037 | MQAKLTKKEFIEWLKTSEGKQFNVDL<br>WYGFQCFDYANAGWKVLFGLLLKGL<br>GAKDIPFANNFDGLATVYQNTPDFLAK<br>PGDMVVFGSNYGAGYGHVAWVIEAT<br>LDYIIVYEQNWLGGGWTDGIEQPGWG<br>WEKVTRRQHAYDFPMWFIRPNFKSET<br>APRSVQSPTQAPKKETAKPQPKAVE | **TKKTSTSNEKWNKNQYGIL<br>WRKEVGSFTCNVPQGIITRR<br>IGPGRQYPIAGALKKAQTV<br>NYTEIQKNDGYIWISWMTN<br>SGYTVYMPVRQVKSDGSLG<br>PLWGTIK** | 99.444 | **65.591** |

**Table S2**. **Sequence source of 8 hits selected after the first screening campaign.**

| ID | Sequence source | |
|---|---|---|
| | EAD | CBD |
| bbst1001 | Literature[1] | SAG |
| bbst1005 | Literature[1] | SAG |
| bbst1014 | Literature[1] | Isolate |
| bbst1021 | SAG | SAG |
| bbst1027 | SAG | SAG |
| bbst1035 | Isolate | Isolate |
| bbst1036 | Isolate | Isolate |
| bbst1037 | Isolate | SAG |

**Table S3 Effects of endolysins against clinical isolates of MRSA**.

The MIC values against nine clinical isolates of MRSA (No.82~99) were determined. The $MIC_{50}$, $MIC_{90}$, and geometric mean of MIC were shown. $MIC_{50}$ and $MIC_{90}$ indicate the MIC values at which the growth of 50% and 90% of the clinical isolates were inhibited, respectively.

| ID | MRSA MIC (μg/mL) (Clinical isolates) | | | | | | | | | | | |
|---|---|---|---|---|---|---|---|---|---|---|---|---|
| | No. 82 | No. 83 | No. 84 | No. 89 | No. 90 | No. 91 | No. 93 | No. 95 | No. 98 | No. 99 | **$MIC_{50}$** | **$MIC_{90}$** |
| bbst1001 | 2 | N.A. | 4 | 8 | 8 | 1 | N.A. | 16 | 4 | N.A. | **4** | **16** |
| bbst1005 | 16 | >32 | 32 | N.A. | N.A. | 8 | N.A. | N.A. | N.A. | 16 | **16** | **>32** |
| bbst1014 | 16 | 8 | 8 | 8 | 16 | 8 | 32 | 8 | >32 | 32 | **8** | **32** |
| bbst1021 | 32 | >32 | 16 | >32 | 4 | 8 | >32 | >32 | N.A. | 16 | **32** | **>32** |
| bbst1027 | 4 | 8 | >32 | 8 | 16 | 8 | >32 | 4 | 8 | 4 | **8** | **>32** |
| bbst1035 | 32 | 8 | 8 | >32 | 16 | 2 | 2 | 8 | 4 | 16 | **8** | **>32** |
| bbst1036 | 1 | 8 | 8 | 2 | 4 | 1 | 2 | 1 | 0.5 | 2 | **2** | **8** |
| bbst1037 | 32 | 2 | 8 | >32 | 8 | 2 | 4 | 2 | 8 | 8 | **8** | **32** |
| Vancomycin | 4 | 2 | 1 | 2 | 1 | 1 | 4 | 1 | 1 | 4 | **1** | **4** |
| Oxacillin | 8 | >32 | >32 | 32 | >32 | >32 | 32 | >32 | >32 | >32 | **>32** | **>32** |

**Hemolytic toxicity assay**

The human whole blood (COSMO Bio) was centrifuged at 800$g$ for 10 minutes and washed with PBS three times. And then, diluted ten-fold with PBS to prepare for 10% human red blood cells (hRBC) solution. The 0.1 mL of test compounds were mixed with an equal volume of 10% hRBC solution, and the reaction mixture was incubated at 37 °C for 1 hour. The 10% hRBC solution was treated with 1% Triton-X100 (Sigma-Aldrich) as well to prepare for complete hemolysis control. After incubation, the reaction mixture was centrifuged at 800$g$ for 10 minutes, and the 20 µL of supernatant was diluted five-fold with PBS. Then the OD405 of each sample was measured. Hemolysis (%) was calculated by the following equation:

$$Hemolysis\ (\%) = \frac{(OD_A - OD_{PBS})}{(OD_{Ctrl} - OD_{PBS})} \times 100$$

,wherein $OD_A$ means $OD_{405nm}$ value of the samples treated with test compounds, $OD_{PBS}$ is that treated with PBS as a no hemolytic condition, $OD_{Ctrl}$ is that treated with Triton-X100.

**Cytotoxicity assay**

The HCT116 cells, which is the human cell line derived from colon cancer, were maintained with McCoy's 5A medium with 10% FBS at 37 °C in 5% $CO_2$. The cells were grown until 80% confluent condition, the cells were collected and distributed in the 96-well tissue culture plate at $3\times10^3$ cells/well. Then, the cells were treated with test compounds at 37 °C in 5% $CO_2$ for 24 hours. After incubation, 10 µL of CCK-8 solution was added to the 110 µL of each cell culture treated with test compounds. And then the mixture was incubated at 37 °C in 5% $CO_2$ for 4 hours, and the absorbance at 450 nm of them was measured to evaluate the cell viability.

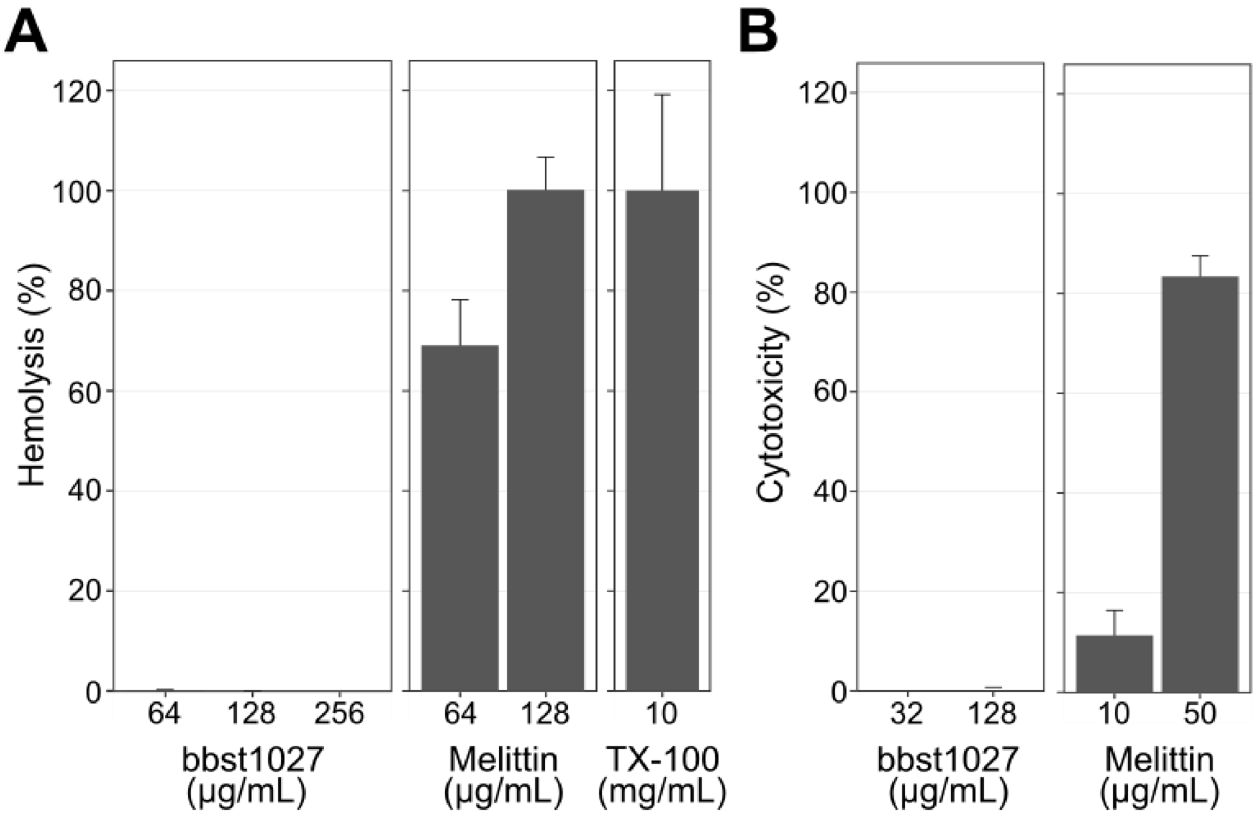


**Fig.S2 Cytotoxicity of bbst1027.** (A) Hemotoxicity of bbst1027 to human blood cells. 1% Triton X-100(TX-100) was used as a control for complete hemolysis. Melittin, which is a peptide derived from bee venom, was used as a positive control. (B) Cytotoxicity of bbst1027 to the colon cancer cell line HCT-116. Melittin was used as a positive control.
